# Inhibition of IL-1β secretion and mitochondria respiration by arsenite which acts on myocardial ischemia-reperfusion injury

**DOI:** 10.1101/751297

**Authors:** Min Li, Yingwu Mei, Jingeng Liu, Kaikai Fan, Xinyue Gu, Jitian Xu, Yuebai Li, Yang Mi

## Abstract

Arsenite (NaAsO_2_) is a potent toxin that significantly contributes to human pathogenesis. Chronic exposure to arsenite results in various diseases. The physiologically important biological target(s) of arsenite exposure is largely unknown. Here we found that transient sodium arsenite treatment (1) blocks nigericin or Rotenone induced IL-1β secretion; (2) inhibits mitochondrial respiration with complex I–linked substrate; (3) induces Heme oxygenase-1 (HO-1) in myocardial tissue. (4) attenuates the myocardial ischemia-reperfusion injury in an in vivo model of rats. The causal relationship among these activities needs further investigation.

## INTRODUCTION

Myocardial ischemia-reperfusion (I/R) injury is an important clinical issue. Great efforts have been devoted to alleviate I/R injury which can greatly improve the patient’s prognosis ^[1]^.

NLRP3 inflammasome mediates the sterile inflammatory response triggered by tissue damage and plays a role in the pathogenesis of myocardial ischemia-reperfusion injury ^[2-4]^. Blocking inflammasome activation by genetic method markedly diminishes infarct development and myocardial fibrosis and dysfunction ^[2-3]^.

Mitochondrial respiration rate is also involved in myocardial I/R injury ^[5]^. Hydrogen sulfide, a potent and reversible inhibitor of cytochrome c oxidase (complex IV of the mitochondrial electron transport chain) has been proven to attenuate myocardial ischemia-reperfusion injury. Possible mechanism for H_2_S protective action on mitochondrial function may lie in its ability to modulate cellular respiration, which limiting the generation of reactive oxygen species and diminishing the degree of mitochondrial uncoupling in perfusion stage ^[6]^. Complex I inhibitors such as rotenone also decrease ROS and oxidative damage during IR and protect against reperfusion injury ^[7-8]^.

In a screening for chemicals that influence inflammasome activation, one hit that drew our attention was the compound sodium arsenite. Arsenic is a metalloid that generates various biological effects on living organism. Acute arsenite toxicity is considered as oxidation of cysteine residues in target proteins that directly alters their conformation or activity ^[9]^. High dose arsenite is reported to block the oxidation of α-oxoglutarate ^[10]^. Cells exposed to arsenite start a cytosolic stress response including phosphorylation of eukaryotic translation initiation factor 2 alpha (eIF2α) and elevation of downstream genes such as CHOP, GADD34 and heat shock proteins which are important against stressful insults ^[11-16]^. Sodium arsenite also induces heme oxygenase-1 (HO-1) which protects against ischemia and reperfusion injury in mice ^[17-19]^.

Considering the pros and cons of arsenite on cells, we wonder its effect on myocardial ischemia-reperfusion injury. The in vivo myocardial ischemia-reperfusion model in rats was made to explore the effect of arsenite on myocardial tissue during the first 24 hours.

## MATERIALS AND METHODS

### Cell culture and inflammasome stimulation

Human THP-1 cells were grown in RPMI 1640 medium, supplemented with 10% FBS. THP-1 cell were differentiated for 3 hr with 100 nM phorbol-12-myristate-13-acetate (PMA). Bone marrow was flushed from mouse femurs and cultured in bone marrow media (RPMI 1640 containing 20% heat-inactivated fetal bovine serum, 30% L929 cell media, 2 mM L-glutamine) for 7 d at 37°C in 4 100-mm polystyrene dishes to obtain mature, differentiated macrophages.

For inducing IL-1β, 10^6^ macrophages were plated in 12-well plate overnight and the medium was changed to opti-MEM in the following morning, and then the cells were primed with ultrapure LPS (100 ng/ml) for 3 hr. After that, arsenite were added into the culture for 5 minutes, and then the cells were stimulated with ATP (5 mM, pH adjusted to 7.5) or Nigericin (10 μM) for 1 hour, or rotenone (10 μM) for 3 h.

### Immunostaining

U2OS or HeLa cells were plated on coverslips for 1 day and transfected with pcDNA6-flag-hNLRP3 vector for 36 hours. After that, the cells were treated with nigericin (10 μM) for 30min and stained with antibody.

For immunostaining, cells were fixed with 4% paraformaldehyde for 15 minutes and permeabilized with 0.1% Triton X-100 in PBS, before incubation with anti-FLAG M2 antibody followed by Cy3 secondary antibodies. Nuclei were stained with DAPI.

### Isolation of mitochondria

Deprive a 150g to 200g male Sprague-Dawley rat of food overnight. Liver was taken out immediately after animal was sacrificed and washed with ice-cold LHM (0.2M mannitol, 50mM sucrose, 10mM KCl, 100mM HEPES, 1mM EDTA). Then the liver was minced finely and washed with LHM again. The minced tissue was homogenized with Dounce homogenizer and centrifuged at 1000 g at 4°C for 5 min to remove nuclei and tissue debris. The supernatant was transferred and centrifuged at 4000 g at 4°C for 10 min. The pellet suspended with LHM was homogenized 3 to 4 times gently again and centrifuged at 4000 g at 4°C for 10 min. The pellet was washed with LHM again. The resulting pellet was re-suspended in respiration medium.

### Respiration measurements

Respiration measurements in isolated mitochondria were conducted at 25°C in a Clark type oxygen electrode (INESA Scientific Instrument, REX, JPB-607A portable Dissolved Oxygen Meters). The respiration medium consisted of 0.225M mannitol, 55mM sucrose, 10mM KH2PO_4_, 12mM KCl, 5mM MgCl_2_, 10mM Tris-HCl pH 7.4. Mitochondria were transferred to a electrode chamber placed on a magnetic stirrer and the volume was adjusted to 2 ml. Mitochondria were treated with arsenite or not 4 min before oxygen consumption was recorded. The effects of arsenite on mitochondrial respiration were determined in the presence of excess ADP (1 mM), using substrate combinations targeting either Complex I (5 mM glutamate plus 2 mM malate) or Complex II (5mM succinate plus 1.25 μM rotenone) of the respiratory chain.

### In vivo myocardial I/R

SD rats (250-300g) were intraperitoneally anesthetized with sodium Pentobarbital (40mg/kg), orally intubated with Teflon tubes (1.2/1.6mm), and connected to a rodent ventilator. A left thoracotomy was then performed. The chest muscle was set aside, and the third and fourth ribs were break off. The area around left auricle and pulmonary arterial cone was exposed. The left anterior descending (LAD) coronary artery was ligated using 6-0 silk suture around a fine tube with a firm knot. Rats were subjected to 40 minutes of LAD ischemia followed by 24h of reperfusion. The infarct area was determined by 2,3,5-triphenyltetrazolium chloride (TTC) staining. All animal studies were conducted in accordance with the National Institutes of Health (NIH) Guide for the Care and Use of Laboratory Animals. All animal experiments performed in this study adhered to the protocols approved by the Institutional Animal Care and Use Committee of Zhengzhou University.

### Infarct area assessment

After 1-day of reperfusion, rats were anesthetized, Evans blue solution (3%, 1ml) were directly injected into the right ventricle, 1 minute later, the heart was quickly excised, immediately frozen, and sliced. Sections were then incubated in a 1% TTC solution which was freshly prepared, and the slices must be covered with a glass sheet during staining. After staining, sections were digitally photographed and the infarct area was determined by computerized planimetry.

### Echocardiography

After 24 hr of reperfusion, echocardiography was performed using a VEVO 2100 high-resolution in vivo imaging system (FUJIFILM Visual Sonics, Toronto, Ontario, Canada). Under anesthesia, the chest of the rat was shaved, and two-dimensional echocardiography was performed. Cardiac function was evaluated by M-mode. Left ventricular (LV) end-diastolic diameter and LV end-systolic diameter were measured on the parasternal LV long-axis or short-axis view. Left ventricular ejection fraction (EF) and fractional shortening (FS) were calculated with computerized algorithms. All measurements represented the mean of five consecutive cardiac cycles.

### Arterial pressure measurements

After 24 hr of reperfusion, before TTC staining method was performed, mice were anesthetized. The femoral artery was dissected and was cannulated using a catheter pre-filled with heparinized normal saline (0.5 IU/ml). After cannulation, the other end of the catheter was connected to the transducer. The arterial systolic pressure and diastolic pressure was monitored by a biological Signal Acquisition and Analysis System (BL-420F, TECHMAN, Chengdu, China).

### Western blotting

The antibodies for phospho-eIF2 Ser51 were purchased from elabscience (E-AB-20864). The antibodies for eIF2α (sc-133132), GADD34 (sc-373815), GADD153 (sc-7351) and Heme Oxygenase 1 (sc-136960) were from Santa Cruz biotechnology.

Mice intraperitoneally injected with arsenite (4.5ug/g) for various periods were sacrificed and organs were excised quickly. Protein was extracted from frozen tissue by grinding using a pre-chilled mortar. The RIPA lysis buffer (50mM Hepes pH7.5, 150mM NaCl, 2mM EDTA, 2mM EGTA, 1% TritonX-100, 50mM NaF, 5mM Sodium Pyrophosphate, 50mM Sodium glycerophosphate, 1mM NaVO_3_, 1mM DTT, 1mM PMSF, 10g/ml Leupeptin, 10g/ml Aprotinin) was used for protein preparation. Following SDS-PAGE, nitrocellulose membranes were blocked with 3% skim milk, and immunoblotting was performed as described elsewhere.

### RT-PCR

Total RNA was extracted from hearts, brain and liver tissue of rats intraperitoneally injected with PBS or arsenite (4.5ug/g) for 7 hours. Semi-quantitive PCR were performed to detect the mRNA expression. The following primers were used: HO-1 (5’-GACCGTGGCAGTGGGAATT-3’ and 5’- TGGTCAGTCAACATGGACGC-3’), CHOP (5’-ACTCTTGACCCTGCATCCCT-3’ and 5’-TCTCATTCTCCTGCTCCTTCTC-3’), XBP1 (5’-TTAGTGTCTAAAGCCACCCACC-3’ and 5’-GCCAGGCTGAACGATAACTG-3’), GAPDH (5’-TGGAAAGCTGTGGCGTGAT-3’ and 5’-GGGTGGTCCAGGGTTTCTT

### Statistical analysis

Data are presented as mean± standard deviation. Comparisons between groups were performed by Student’s two-tailed unpaired t-test. Statistical significance was set at P <0.05.

## RESULTS

### Arsenite blocks nigericin or Rotenone induced IL-1β secretion

In a screening for chemicals that acted on inflammasome activation, we found that sodium arsenite strongly inhibited IL-1β secretion. To determine the effects of arsenite on NLRP3 inflammasome activation, bone-marrow-derived macrophages (BMDMs) or THP-1 cells were treated with arsenite before LPS priming or before agonist challenge. We observed that arsenite blocked IL-1b secretion at both situations. It has been reported that arsenite can inhibit protein synthesis ^[9]^ and NF-κB activation ^[20]^, we then examined whether arsenite had an impact on LPS-induced priming for inflammasome activation. As shown in Figure 1C, Arsenite inhibited LPS-induced pro-IL-1b, pro-caspase-1 and NLRP3 expression.

**Figure 1.**
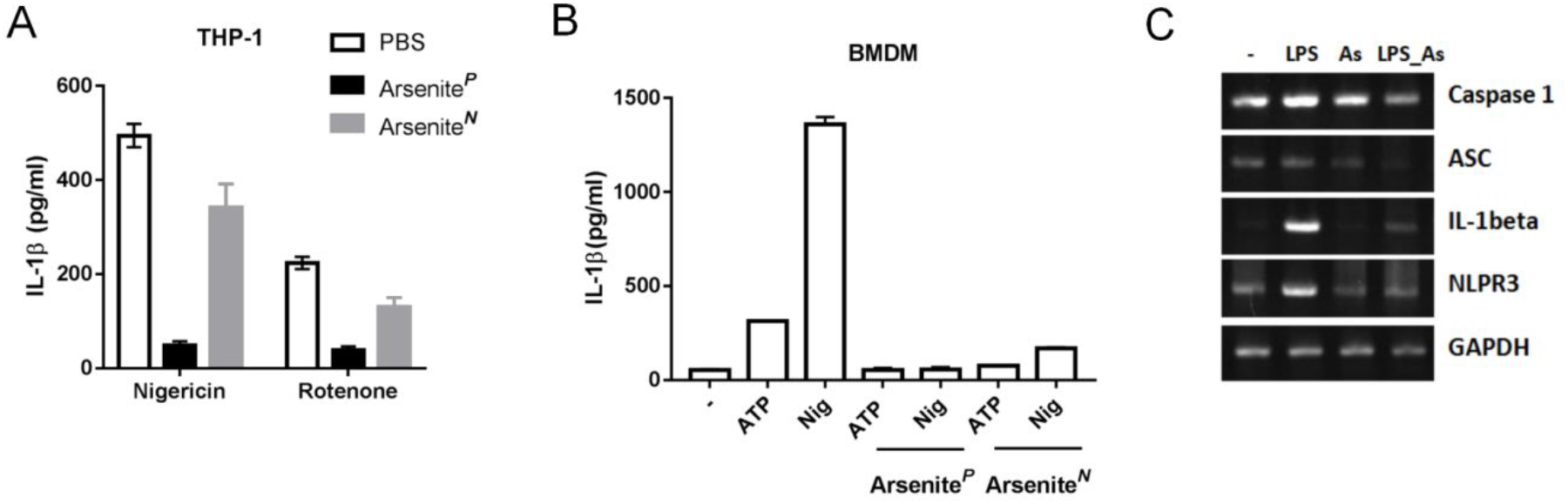
Arsenite blocks NLRP3 inflammasome activation. (A) THP-1 cells were primed with LPS (100 ng/ml) for 3 hr and were stimulated with nigericin (5μM) for 1 hour or Rotenone (10μM) for 3 hours. Sodium arsenite (40μM) was added into medium together with LPS (Arsenite^*P*^) or with Nigericin (Arsenite^*N*^). Supernatants were analyzed by ELISA for IL-1β. Experiments were repeated three times. (B) LPS-primed bone-marrow-derived macrophages (BMDMs) were stimulated for 1 hour with ATP (5 mM) or Nigericin (10μM), Sodium arsenite (10μM) was added into medium together with LPS (Arsenite^*P*^) or with Nigericin (Arsenite^*N*^). Supernatants were analyzed by ELISA for IL-1β. Experiments were repeated three times. (C) BMDMs treated with LPS and arsenite alone or collectively for 4 hours were analyzed by RT-PCR. “As”, Sodium arsenite.

**Figure 2.**
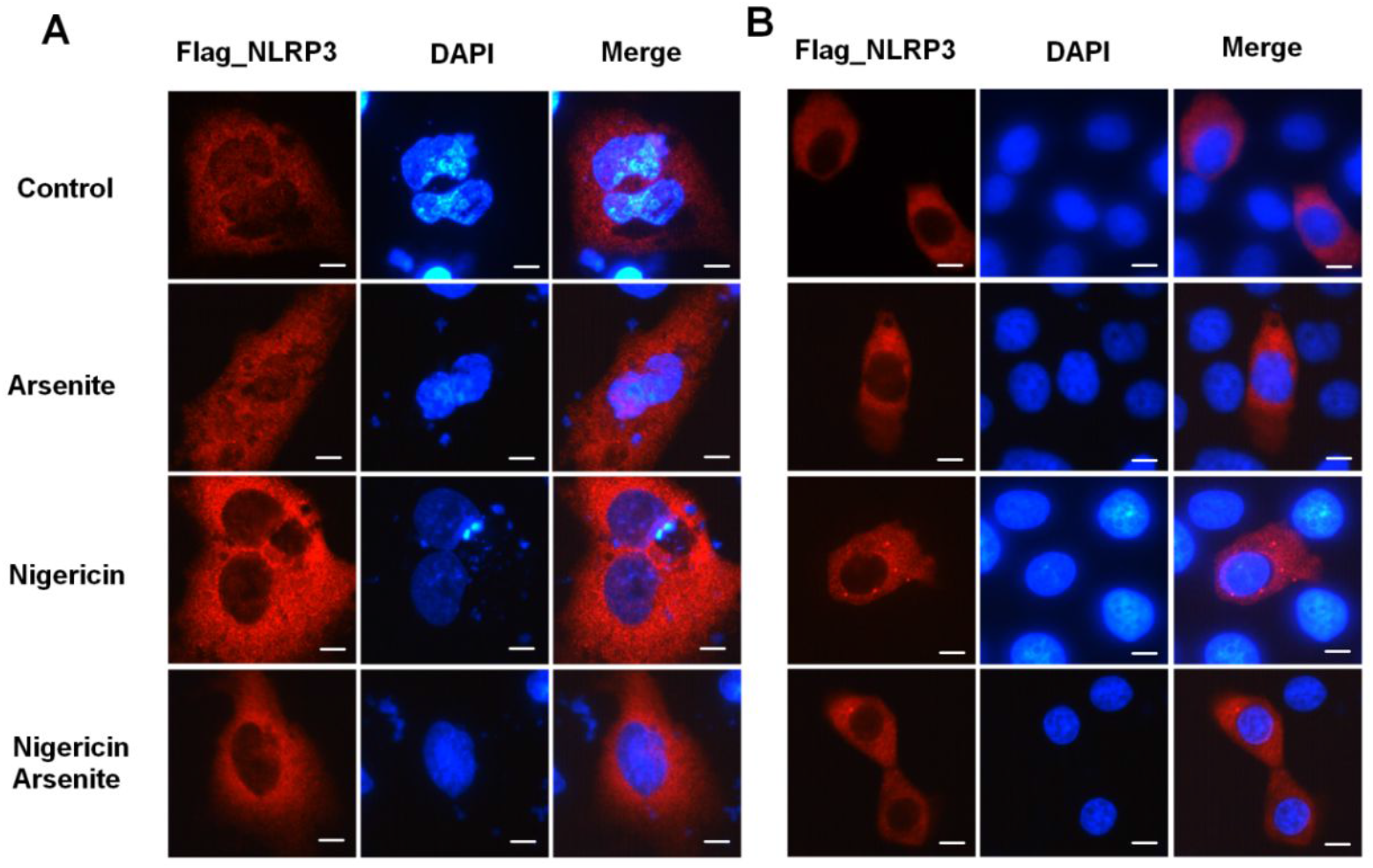
NLRP3 aggregation is compromised by arsenite. (A) U2OS and (B) HeLa cells transfected with Flag_NLRP3 vector were treated with nigericin and sodium arsenite alone or collectively for 30 minutes, cells were fixed and stained with FLAG antibody. Scale bars: 10μM

It’s recently reported that recruitment of NLRP3 to dispersed trans-Golgi network (dTGN) is an early and common cellular event that leads to NLRP3 aggregation and activation ^[21,22]^. Different NLRP3 stimuli lead to disassembly of the TGN. To explore whether arsenite has any effect on the recruitment of NLRP3 to dTGN, U2OS and HeLa cells transfected with flag tagged NLRP3 were treated first with arsenite and then with nigericin. Fluorescence microscopy experiments showed that flag_NLRP3 aggregated around nucleus in U2OS or formed small dots in HeLa cells upon nigericin treatment, whereas arsenite decreased the NLRP3 aggregation.

### Inhibition of mitochondrial respiration by high dose arsenite

Arsenite was reported to have an effect on mitochondria. We explored its role on respiration rate. Arsenite at 200uM produce a strong inhibition of oxygen consumption in isolated rat liver mitochondria with complex I–linked substrate (Figure 3B, 3C), but did not have a substantial effect on complex II–linked respiration using succinate as the substrate (Figure 3A). Arsenite at 20uM did not show a significant effect on complex I (Figure 3C).

**Figure 3.**
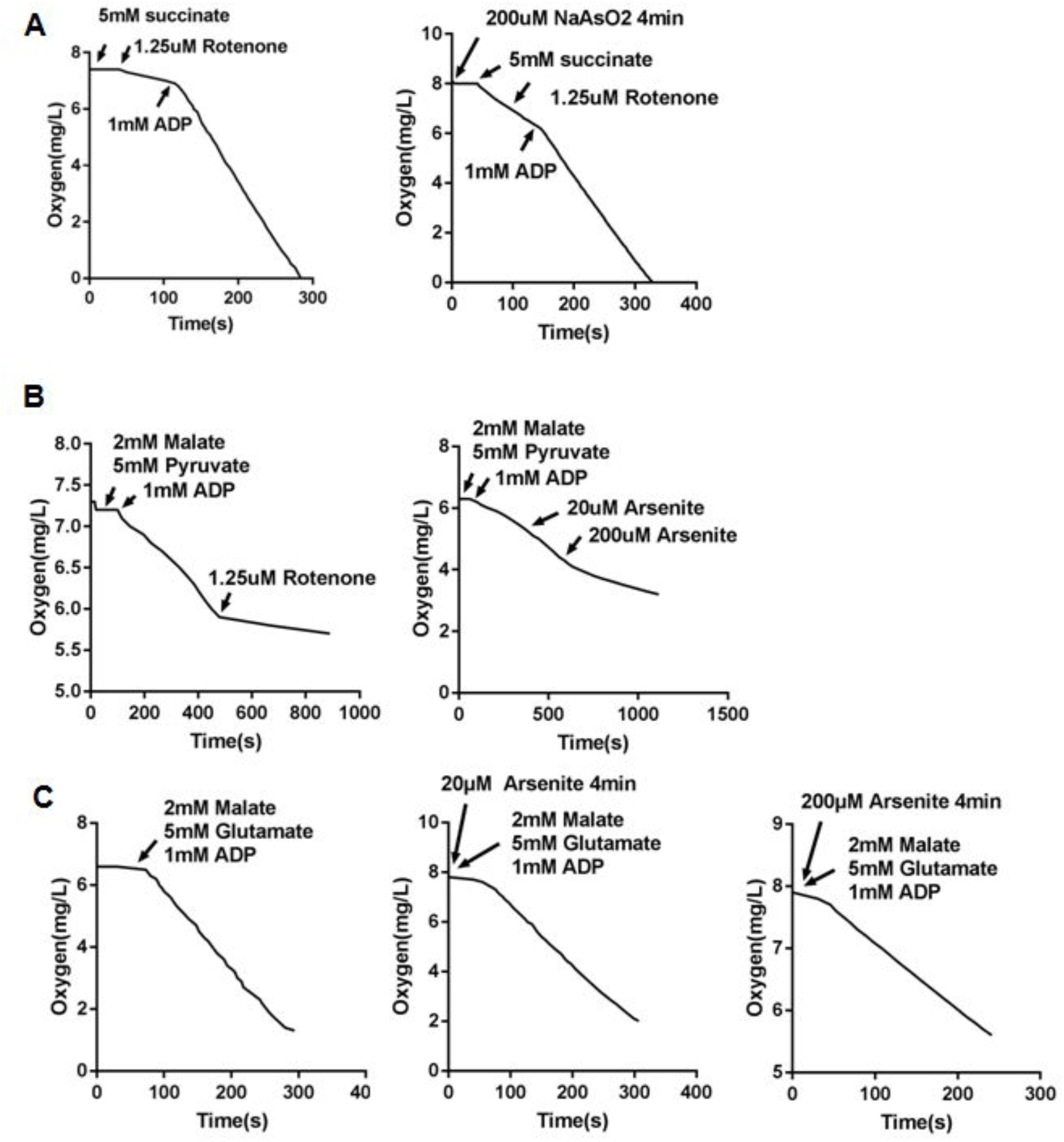
Inhibition of mitochondrial respiration in isolated rat liver mitochondria by sodium arsenite. (A) Mitochondria were treated with sodium arsenite for 4min or not, and then oxygen consumption rates were measured using substrate (5 mM succinate plus 1.25μM rotenone) targeting respiratory complex II (B) Oxygen consumption rates were measured using substrate (2 mM malate plus 5 mM Pyruvate) targeting respiratory complex I. Rotenone or NaAsO_2_ were added into the reaction buffer as the arrows showed. (C) Mitochondria were treated with sodium arsenite for 4min or not, and then oxygen consumption rates were measured using substrate (2 mM malate plus 5mM glutamate) targeting respiratory complex I.

**Figure 4.**
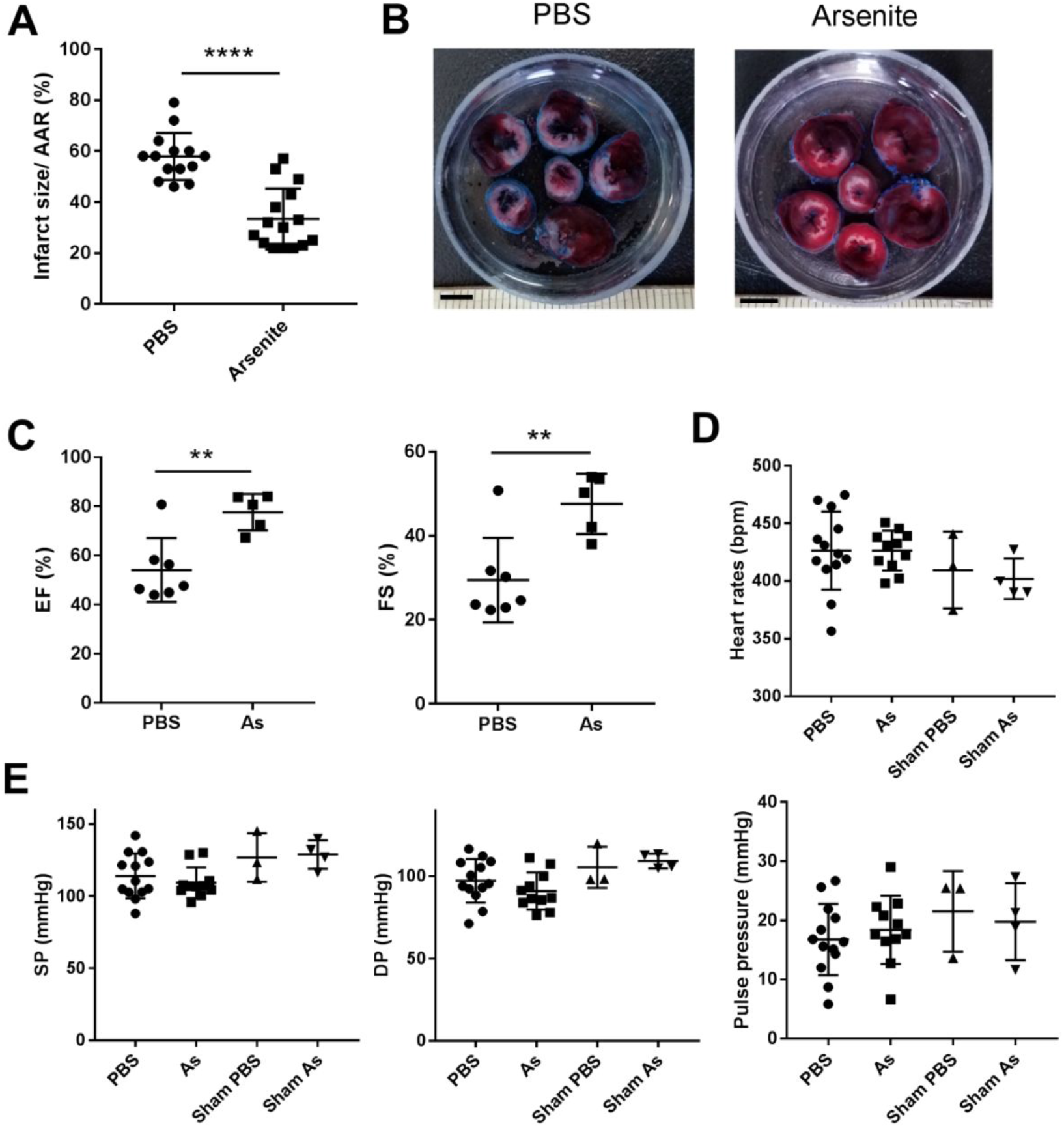
Effect of arsenite treatment on myocardial I/R. (A) Quantification of infarct size of myocardial tissues 1 day after reperfusion (n=14 and 15) by TTC staining method. Rats were intraperitoneally administrated with PBS or 4.5mg/kg arsenite 30min before LCA was occluded. The data were analyzed using t-test, P<0.0001. AAR, the area at risk. (B) Representative images of heart slices from different groups at 1 day after reperfusion. The non-ischemic area is indicated in blue, the area at risk in red, and the infarct area in white. Scale bars: 5mM (C) Ejection fraction (EF, %) and LV fractional shortening (FS, %) (n =5-7) analyzed by M-mode images of the LV from PBS and arsenite group 1 day after myocardial I/R. The data were analyzed using t-test, ** P< 0.01. As, Sodium Arsenite. (D) and (E) The heart rates, systolic pressure (SP) and diastolic pressure (DP) were measured by placing a catheter into the femoral artery of rats at 24 hour after myocardial I/R. The pulse pressure was calculated by SP and DP. There is no significant statistic difference of these parameters in each group. As, Sodium Arsenite.

**Figure 5.**
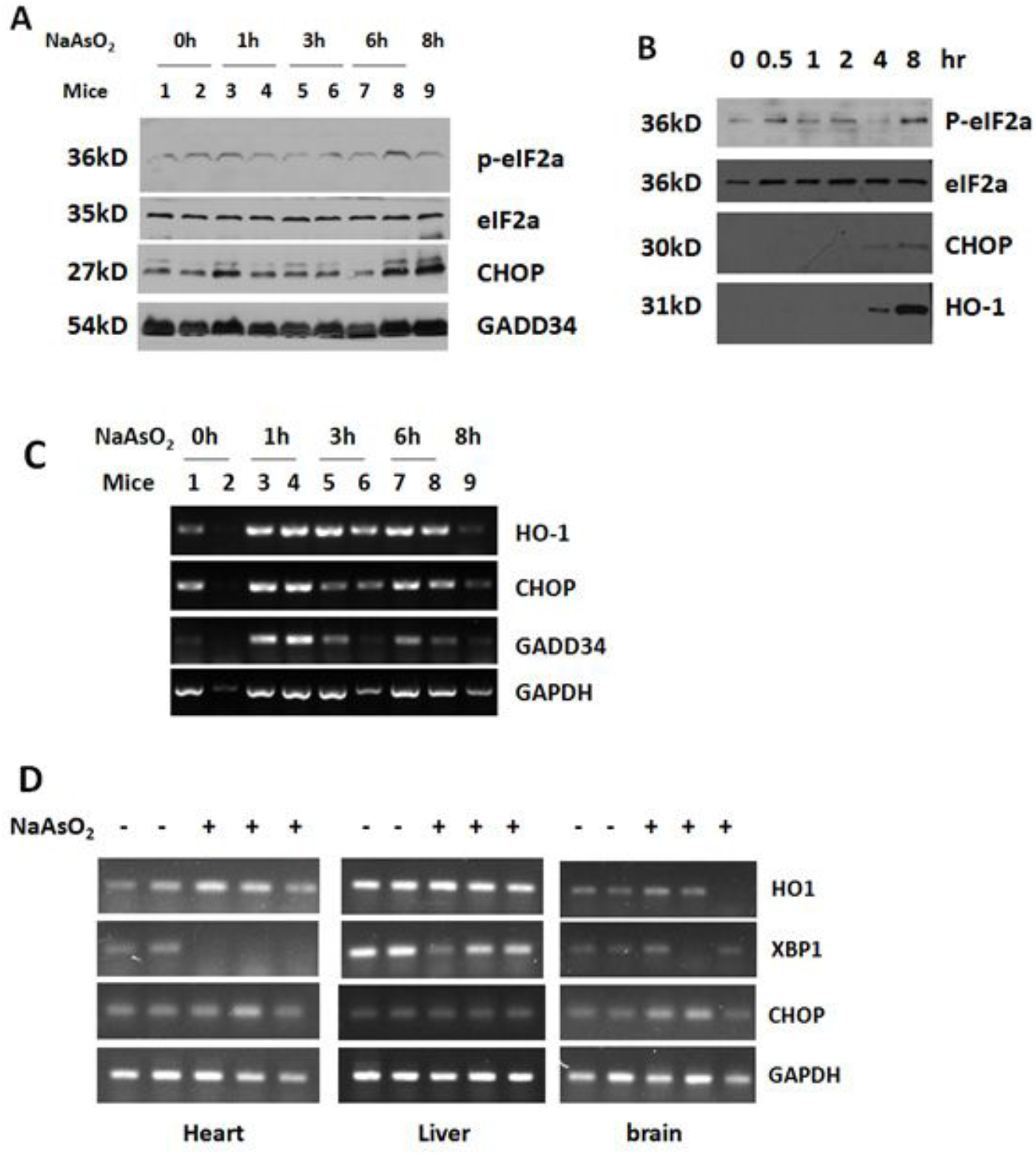
Arsenite induces a transient stress following sustained heme oxygenase-1 expression in myocardial tissue. (A) Western blot analysis of total protein extracted from heart tissue of mice injected with 4.5mg /kg sodium arsenite for the indicated time. (B) Western blot analysis of total protein extracted from HeLa cells treated with 4.5μg /ml NaAsO_2_ for the indicated time. (C) RT-PCR analysis of indicated genes in SMMC-7721cells treated with NaAsO_2_ for the indicated times. (D) SD rats were intraperitoneally injected with 4.5mg /kg NaAsO_2_ for 7 hours. After that total RNA was isolated from heart, liver and brain tissues and stress related genes were analyzed by RT-PCR.

### Arsenite attenuates myocardial ischemia-reperfusion injury

It was reported arsenic trioxide induced apoptosis in cancer cells and here we found that arsenite blocks NLRP3 inflammasome activation which is essential for myocardial ischemia/reperfusion injury, thus we wonder what effect arsenite has on myocardial I/R injury. The in vivo model of myocardial ischemia-reperfusion was used to answer this question. Rats were subjected to 40min of LV ischemia and 24h reperfusion. Arsenite was administered by intraperitoneal injection 30min before ligation of the left anterior descending (LAD) coronary artery at dose of 4.5mg/kg. Evaluation of infarct size revealed a significant cyto-protection effect of arsenite as assessed by 2,3,5-triphenyltetrazolium chloride (TTC) staining. Surprisingly, rats receiving 4.5mg/kg arsenite displayed a lower ratio of infarct size to area-at-risk (INF/AAR), as shown in Figure 2 A and B.

To determine the effect of arsenite on cardiac function after I/R, we measured ejection fraction (EF) and fractional shortening (FS) by echocardiography. The EF and FS were at a relative low level at 1day after myocardial I/R in contrast with the reference, however, treatment with arsenite increased EF and FS after 1 day reperfusion, indicating improved cardiac function. To test whether arsenite affects hemodynamic properties, we measured systolic pressure and diastolic pressure by placing a catheter into the femoral artery of rats at 24 hour after myocardial I/R. The heart rates, systolic pressure, diastolic pressure and pulse pressure were not significantly changed in each group.

### Arsenite induces a transient stress following sustained heme oxygenase-1 expression in myocardial tissue

Exposure to arsenite was shown to induce significant cellular stress, which is manifested as phosphorylation of eIF2α at Ser-51 and elevated expression of heat shock proteins, CHOP and GADD34^[10-17]^.

To explore the biological effects of arsenite on heart, cells or mice were simply treated with arsenite. Arsenite induced phosphorylation of eIF2α in rodent myocardial tissue and HeLa cells as early as in 1h (Fig. 3A, B), which represented a cytosolic stress response is initiated. The mRNA level of stress related proteins such as CHOP and GADD34 was up-regulated transiently in heart tissue at 1h after peritoneal injection of arsenite (Fig. 3B, C). HO-1, which protects cells from oxidative injury, was induced continuously at transcription level by arsenite in heart tissue but not in brain and liver tissue. HO-1 protein expression can be seen as early as 4 hours after arsenite treatment.

## DISCUSSION

The major findings of this study are as follows: (1) Arsenite blocks nigericin or Rotenone induced IL-1β secretion by inhibiting both the priming and activation stage; (2) Mitochondrial complex I respiration is inhibited by high dose arsenite; (3) Arsenite attenuates the myocardial ischemia-reperfusion injury in rats; (4) Arsenite induces HO-1 in myocardial tissue. However, the causal relationship among these activities needs further proof. The relationship between NLRP3 inflammasome and ischemia-reperfusion injury has been studied for several years. Arsenite at least partially acts on myocardial tissue by this way.

Arsenite at 200uM strongly inhibits oxygen consumption in isolated rat liver mitochondria with complex I–linked substrate. Whether arsenite targets the complex I or related dehydrogenates needs further investigation. Complex I is the entry point for electrons from NADH into the mitochondrial respiratory chain and is well established as a major source of mitochondrial superoxide in vitro ^[5]^. There is evidence that complex I is the major site of mitochondrial superoxide production upon reperfusion ^[5, 7-8]^. So inhibition of complex I will decrease ROS and oxidative damage during IR. Respiration inhibition may also decrease the overall metabolic rates and save nutrients and oxygen consumption. Arsenite at 20uM does not show significant inhibitory activity on respiration of isolated mitochondria.

Arsenite induces a cytosolic stress characterized by eIF2α phosphorylation and downstream events before myocardial ischemia. Pre- and post conditioning with ischemia are representative strategies to reduce infarct size in animal models ^[23, 24]^. Both of these treatments in fact induce a cellular stress response, and arsenite may induce a similar cardiac stress response which is protective for the subsequent ischemia injury.

Heme oxygenase-1 and heat shock protein induction may contribute for cell survival during I/R. By degrading the oxidant heme and generating the antioxidant bilirubin and anti-inflammatory molecule carbon monoxide (CO), HO-1 protects cells from death due to path-physiological stress and oxidative injury. HO-1 has been shown to play a role in limiting myocardial Ischemia/Reperfusion injury in HO-1 knockout or transgenic mice. HO-1 protein level can be seen increased in 4 hours after arsenite treatment.

## ABBREVIATIONS

HO-1: Heme oxygenase-1
I/R: ischemia/reperfusion
LV: left ventricular
eIF2α: eukaryotic translation initiation factor 2 alpha
As_2_O_3_: Arsenic trioxide
APL: acute promyelocytic leukemia
LAD: The left anterior descending
TTC: 2,3,5-triphenyltetrazolium chloride
EF: ejection fraction
FS: fractional shortening
PMA: phorbol-12-myristate-13-acetate
BMDM: bone-marrow-derived macrophages
dTGN: dispersed trans-Golgi network
INF/AAR: infarct size to area-at-risk.

## ACKNOWLEDGEMENTS

We thank Dr Shengna Han and Shuhui Wang from Department of Pharmacology of Zhengzhou University for their help with establishing in vivo heart IR model. This work was supported by “Youth Initiation Fund of Zhengzhou University [1411328011]”.

## AUTHOR CONTRIBUTIONS

Min Li performed most of the experiments. Yang Mi designed the whole project including every experiment, analyzed all the data, performed some experiments and prepared the manuscript. PhD Yingwu Mei, Professor Jitian Xu and Professor Yuebai Li kindly provide some materials and advices of some experiments. PhD Jingeng Liu and PhD Kaikai Fan help to establish the myocardial I/R model.

## CONFLICT OF INTEREST

The authors declare that they have no conflict of interest.

## Notes

#### Summary of Updates

The title, abstract and discussion were revised; New data (Fig.2 and Fig.3) added.

